# Probing the ligand binding specificity of FNBP4 WW domains and interaction with FH1 domain of FMN1

**DOI:** 10.1101/2023.02.13.528256

**Authors:** Shubham Das, Sankar Maiti

**Affiliations:** Department of Biological Sciences, Indian Institute of Science Education and Research Kolkata, Mohanpur, Nadia - 741246, West Bengal, India

**Keywords:** FNBP4, FMN1, FH1 domain, WW domain, Protein-protein interaction, Surface plasmon resonance

## Abstract

Formins are a group of actin-binding proteins that mediate nascent actin filament polymerization, filament elongation, and barbed end capping function, thereby regulating different cellular and developmental processes. Developmental processes like vertebrate gastrulation, neural growth cone dynamics, and limb development require formins to function in a regulated manner. Formin binding proteins like Rho GTPase regulates activation of auto-inhibited conformation of diaphanous formins. Unlike other diaphanous formins, Formin1 (FMN1), a non-diaphanous formin, is not regulated by Rho GTPase. FMN1 acts as an antagonist of the BMP signaling pathway during limb development. Several previous reports demonstrated that WW domain-containing proteins can interact with poly-proline-rich amino acid stretches of formins and play a crucial role in developmental processes. WW domain-containing FNBP4 protein plays an essential role in limb development. It has been hypothesized that the interaction between FNBP4 and FMN1 can further attribute to the role in limb development through the BMP signaling pathway. In this study, we have elucidated the binding kinetics of FNBP4 and FMN1 using surface plasmon resonance and enzyme-linked immunosorbent assays. Our findings confirm that the FNBP4 exhibits interaction with the poly-proline-rich formin homology 1 (FH1) domain of FMN1. Furthermore, only the first WW1 domains is involved in the interaction between the two domains. Thus, this study sheds light on the binding potentialities of WW domains of FNBP4 and their possible contribution to the regulation of FMN1 function.

## 1. INTRODUCTION

Formins have emerged as a multi-domain, sterling group of actin-binding proteins and act as a key mechanistic regulator of actin dynamics.^1,2^ Formins can nucleate nascent actin filament polymerization in vitro by probity of their formin homology 2 (FH2) domain.^3^ Additionally, formins aid filament elongation in processive motion and barbed end capping function.^3^

Formins play a crucial role in developmental processes. For instance, Daam1 (Dishevelled associated activator of morphogenesis 1), a diaphanous related formin regulates the non-canonical planar cell polarity pathway in cell and tissue morphogenesis during gastrulation.^4,5^ Formin 2 (FMN2) plays a role in membrane protrusion in the developing neurons and oocytes.^6^ During renal and limb development, alternatively spliced transcripts of limb deformity genes or formins are predominantly expressed.^7^ Previous reports demonstrated that during limb development the expression of the msh homeobox1 (MSX1) gene and fibroblast growth factor 4 (FGF4) gene at the apical-ectodermal ridge (AER) were regulated by formin1 (FMN1).^8^ FMN1 transcriptionally regulates the cis-positional neighboring gene Germlin expression on chromosome 2.^9^ Germlin acts as an antagonist of bone morphogenetic protein (BMP), whereas FMN1 functions as a repressor of Smad phosphorylation in the BMP signaling pathway. So Germlin and FMN1 both are acting as negative regulators of the BMP signaling pathway.^8,10^

Regulation of formins is essential for the above-mentioned developmental processes. Proteins like Rho GTPase, an essential group of formin-binding proteins (FBP), release an auto-inhibited conformation of diaphanous formins.^11–13^ Non-diaphanous formins like FMN1, INF, Delphilin, etc. are not regulated by Rho GTPase.^1,2^ There is still a dearth of knowledge about the regulatory mechanism of non-diaphanous formins. Further study is required to understand the mechanisms perspicuously.

Numerous proteins with SH3 and/or WW domains can bind to formins; classified as Formin-binding proteins (FBPs).^14,15^ WW domains are comprised of 38 to 40 amino acid residues and nomenclature is based on presence of two tryptophan (W) amino acid; having an anti-parallel triple-stranded beta-sheet and linked with two beta turns.^15,16^ Proline-rich sequences are prevalent in the inter-domain region of multi domain containing proteins and this poly-proline-rich sequences can bind to WW domain.^17^ WW domains appear in tandem repeats in several multiple WW domain-containing proteins. Notably, FBPs like NEDD-4, YAP-65, FBP11, FBP21, FNBP4, etc contain multiple WW domains.^15^ Despite this, information regarding the need for the presence of these multiple copies of WW domains in different proteins or regulations of such tandem WW domains is not well studied. Though several proteins with WW domains have been categorized into one group, these proteins play distinct roles in cells. For example, NEDD-4 helps in differentiation of the central nervous system.^18^ During cancer cell migration FBP17 helps in extracellular matrix degradation and invadopodia formation.^19^ YAP-65 acts as a transcription regulator through chromatin remodeling.^20^ Interestingly, FNBP4 (also known as FBP30) plays an essential role in eye and limb development. Chan *et al*. reported the mRNA expression of FBPs like FBP17, FBP27, FBP30, etc. in 10.5 embryonic day mice.^14^ According to Kondo *et al*., family suffering from microphthalmia with limb anomalies (MLA) has a homozygous mutation in the WW1 domain of FNBP4.^21^

FNBP4 has been annotated as a formin-binding protein but its interaction with FMN1 is yet to be characterized in detail. Characterizing FNBP4 and FMN1 binding interaction led us to elucidate the physiological role of these two proteins. In this present study, we used surface plasmon resonance (SPR) technique and enzyme-linked immunosorbent assay (ELISA) to characterize the binding of FNBP4 and FMN1. We observed that FNBP4 exhibits interactions towards poly-proline-rich FH1 domain of FMN1 but does not interact with the FH2 domain of FMN1. We also studied WW1 and WW2 to collate their binding potentialities and likeliness. In order to demonstrate that we accomplished the binding experiment, the poly-proline-rich FH1 domain of FMN1 interacts with WW1 but does not exhibit interaction with the WW2 domain of FNBP4.

## 2. METHODS

### 2.1. Plasmid construction

C-terminal FH1-FH2 (amino acids 870-1466), C-terminal FH2 (amino acids 983-1466), only FH1 (amino acids 870-970) regions of FMN1 were constructed with pET28a (+) (Novagen). N-terminal WW1-WW2 FNBP4 (amino acids 214-629), WW1 FNBP4 (amino acids 214-430), WW2 FNBP4 (amino acids 384-629), and N-terminal ΔWW1 FNBP4 (amino acids 249-629) were constructed into the vector pET28a (+).

### 2.2. Protein purifications

Plasmid constructs were transformed into *Escherichia coli* BL21 DE3 (Stratagene) stains. Cells were grown in LB media with 30μg/ml kanamycin concentration at 37 °C and induced at 0.5 O.D. at A600, with 0.5 mM IPTG (Himedia) at 19 °C for 12 hours (For FNBP4 constructs were induced at 25 °C for 8 hours). Cells were harvested then resuspended and lysed by sonication in lysis buffer (0.2% IGEPAL, 150 mM NaCl, 30 mM imidazole at pH 8, 0.5 mM DTT, 50 mM Tris at pH 8) with a protease inhibitor cocktail (Aprotinin, PepstatinA, Leupeptin, Benzamidine hydrochloride, Phenylmethylsulfonyl). Then proteins were affinity purified using Ni^2+^-NTA bead.^22^ The proteins were then eluted with elution buffer (100 mM NaCl, 350 mM imidazole at pH 8, 50 mM Tris at pH 8). For SPR experiment eluted proteins were dialyzed against HBS-N buffer (HEPES-0.01M, NaCl-0.15M pH-7.4) for 4 h at 4 °C.

### 2.3. Antibody production

Anti-FMN1 sera was raised against 6xHis-tagged FH1-FH2 FMN1. BALB/C mice were used for anti-FMN1 sera production using a standard 70 days protocol. Protocol reference no. was IISERK/IAEC/2022/024 has been approved by the institutional animal ethics committee (IAEC). Terminal bleeds for anti-FMN1 antibody checked against recombinant protein by western blot analysis (Fig. S. 1).

### 2.4. Enzyme-linked immunosorbent assay (ELISA)

FMN1 and FNBP4 interaction tested with Enzyme-linked immunosorbent assay (ELISA). 10μg purified FNBP4 was coated into the wells (maxisorp surface) of ELISA plates (#439454 Nunc Maxisorp) along with 1X PBS for the control experiment and allowed to incubate overnight at 4 °C. The wells were blocked with 5% BSA in 1X PBS for 2 hours at room temperature. The wells were then incubated with either FH1-FH2 FMN1 or FH2 FMN1protein for 2 hours at room temperature in a concentration-dependent manner. The bound FH1-FH2 FMN1 and FH2 FMN1 was detected by its specific polyclonal antibodies (raised in mice) at a dilution of 1:1000 Mice anti-FMN1 antibody can detect FNBP4 and FH1-FH2 FMN1 complex or FNBP4 and FH2 FMN1 complex. Followed by HRP-conjugated goat anti-Mouse IgG secondary antibody (#31430 Invitrogen) was added at a dilution of 1:10000 for 45 minutes at room temperature. Color was developed with tetramethylbenzidine (#T0440 Sigma Aldrich) at a 1X dilution for 15-20mins and the reaction ended with H2SO4 (5N). Wells were washed three times with PBST containing 0.02% (v/v) Tween-20 (#M147 Ambrescopropure) in 1X PBS after each incubation step. The absorbance was taken at 450 nm using a microplate reader (Epoch2). Graphs were plotted using Graphpadprism-8.

### 2.6. Surface Plasmon Resonance

The binding kinetics of FNBP4 and FMN1 was determined by SPR BIAcore T200 (GE Healthcare Life sciences). The surface of the CM5 sensor chip (Series S) was activated by EDC/NHS mixture using an amine coupling kit (BR-1000-50, GE Health care Life sciences). 10μg/mL FNBP4 diluted in sodium acetate buffer pH-4.5 (for WW1-FNBP4 fragment pH-4 was used). FNBP4 was immobilized on a CM5 sensor chip in HBS-EP (HEPES-0.01M, EDTA 0.03M, NaCl-0.15M, surfactant P20-0.05%, pH-7.4) running buffer at a flow rate of 30μL/min. After immobilization with FNBP4, ethanolamine was used to block any remaining reactive succinimide ester groups on sensor surface. The non-immobilized flow cell served as reference surface for blank correction. All immobilization experiments were carried out at 25 °C. Different concentrations of analytes were prepared in HBS-EP buffer and flowed over the sensor chip at 30μL/min. HBS-EP buffer was also used as a running buffer. Duplicate concentration for a single concentration of analyte and zero concentration (HBS-EP buffer) were used as positive and negative controls respectively. The association phase was monitored for 120 sec or 180 sec and the dissociation phase was monitored for 300 sec. Regeneration was initiated by 50 mM NaOH solution (GE Healthcare). Obtained SPR sensorgrams were fitted with a 1:1 Langmuir binding model using BIAcore evaluation software, version 2.0. We have also calculated equilibrium dissociation constants (K_D_), association rate (k_a_), and dissociation rate (k_d_).

## 3. RESULT

### 3.1. Proline-rich FH1 domain interacts with WW domains of FNBP4

In order to assess FNBP4 and FMN1 interaction, the binding potentialities of WW1-WW2 FNBP4 to FH1-FH2 FMN1 or FH2 FMN1 had been analyzed, using ELISA and SPR analysis. Based on the domain orientation of FNBP4 and FMN1; WW1-WW2 FNBP4, FH1-FH2 FMN1, and FH2 FMN1 clone constructs were prepared (Fig. 1 A & B). These recombinant WW1-WW2 FNBP4, FH1-FH2 FMN1, and FH2 FMN1fragments were purified as 6x-His tagged with molecular weights of 48.6 kDa, 68.6 kDa, and 56 kDa respectively (Fig. 1 C & D). The functional properties of FH1-FH2 FMN1 and FH2 FMN1 protein fragments had been confirmed with fluorometric pyrene-actin polymerization assays.^23^ Both the protein fragments were active as they had the ability to initiate actin filament polymerization in vitro and induced actin filament polymerization in a concentration gradient manner (Fig. S. 2).

**Fig. 1.**
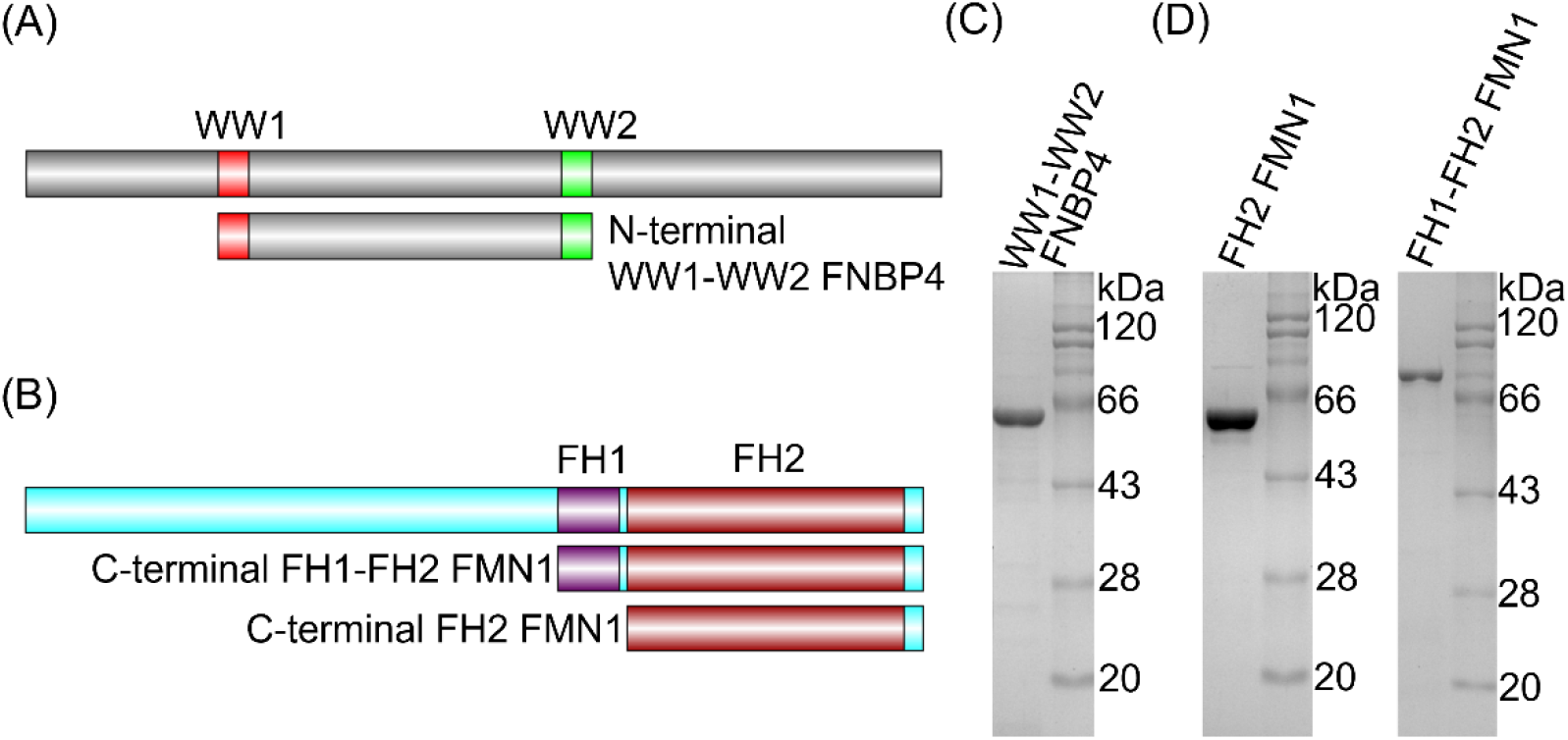
Schematics of constructs and purified fragments of FNBP4 and FMN1. (A) Schematic illustration of N-terminal WW1-WW2 FNBP4 (amino acids 214-629). (B) Schematic representation of FH1-FH2 FMN1 (C-terminal of FMN1: amino acids 870-1466) and FH2 FMN1 (amino acids 983-1466) constructs. (C) Coomassie stained 10% SDS-PAGE of purified 6x-His tagged WW1-WW2 FNBP4 (amino acids 214-629) (48.6 kDa). (D) Purified FH2 FMN1 (amino acids 983-1466) and FH1-FH2 FMN1 (amino acids 870-1466) in Coomassie stained 10% SDS-PAGE (56 kDa and 68.6 kDa respectively).

ELISA-based binding studies were performed where WW1-WW2 FNBP4 was coated in the 96 well plate as a ligand and incubated with different concentrations of FH1-FH2 FMN1 or FH2 FMN1 as analytes.^24^ The absorbance was increased with increasing concentration of FH1-FH2 FMN1 which indicates complex formation between WW1-WW2 FNBP4 and FH1-FH2 FMN1 and reached a saturation of binding (Fig. 2 A). When we used FH2 FMN1 as an analyte there was very negligible absorbance, which indicated an inability for binding to WW1-WW2 FNBP4 (Fig. 2 A).

**Fig. 2.**
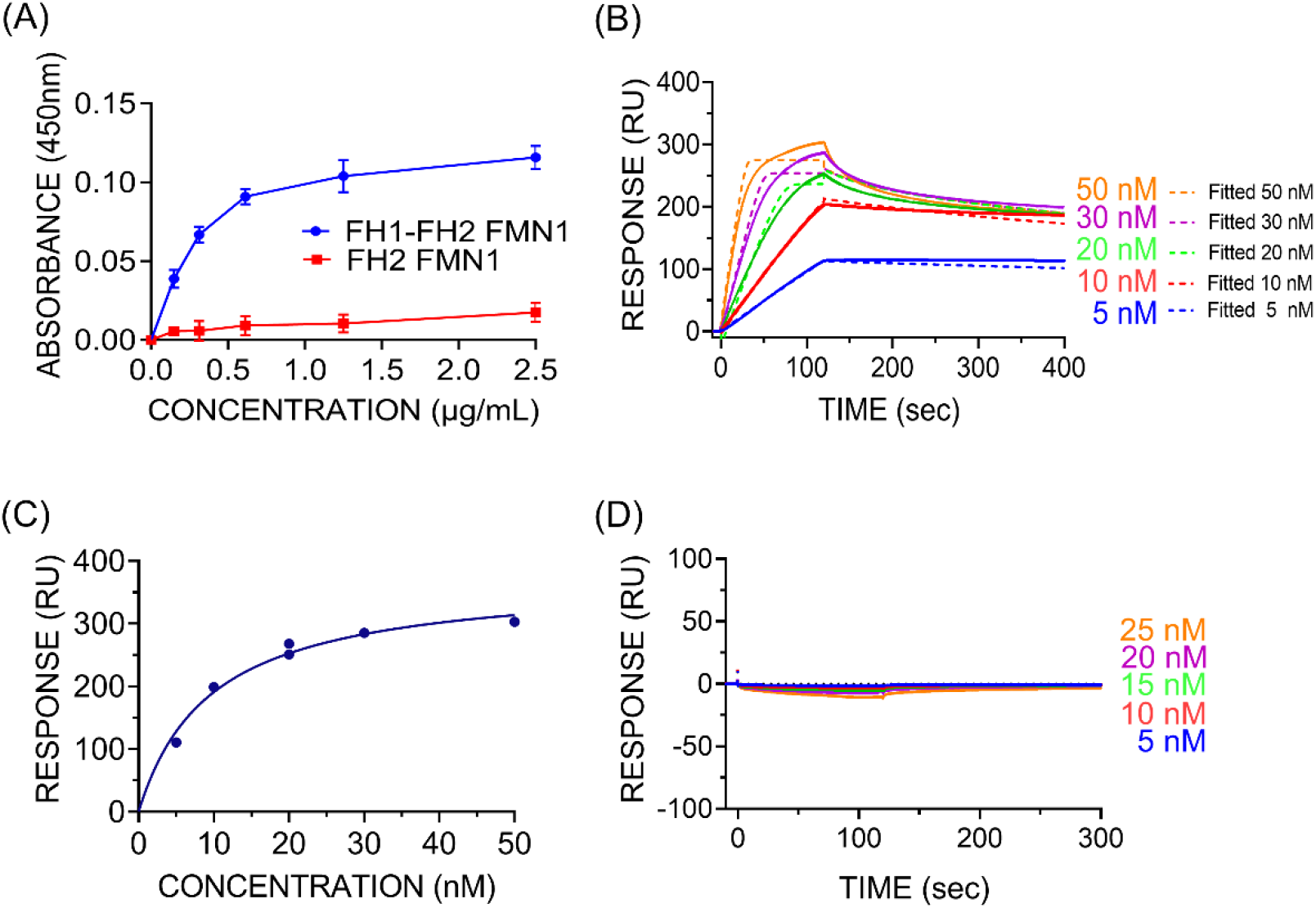
Binding between WW1-WW2 FNBP4 and FMN1. (A) Results of WW1-WW2 FNBP4 and FH1-FH2 FMN1 or FH2 FMN1 interaction obtained by ELISA. WW1-WW2 FNBP4 coated in well. FH1-FH2 FMN1 and FH2 FMN1 were used as analytes. Then mice anti FMN1 Ab (1:1000) was used, followed by anti-IgG HRP conjugated Ab (1:10000) was added. TMB was used as a substrate. Absorbance was taken at 450 nm. (B) SPR sensorgrams of the binding kinetics for WW1-WW2 FNBP4 with FH1-FH2 FMN1, (D) for FH2 FMN1 at 25°C. WW1-WW2 FNBP4 was covalently immobilized on a CM5 (Series S) biosensor surface at pH 4.5 Sensorgrams were plotted as response unit (RU) versus time (Second). Color-coded sensorgrams indicate increasing concentrations of analytes. Sensorgrams were fitted to Langumir binding rate equation and indicated with respective colored dash-line. Sensorgrams were blank-corrected for all cycles. RU depicts, one picogram of analytes bound on a ligand immobilized one mm square surface. (C) Binding affinity curve for FH1-FH2 FMN1 vs WW1-WW2 FNBP4 was fitted with a non-linear regression equation (one-site specific binding model). Duplicate concentration of 20 nM was used for positive control.

Subsequently, SPR analysis had been performed for FH1-FH2 FMN1 with WW1-WW2 FNBP4 to characterize binding kinetics like equilibrium dissociation constant (K_D_), association constant (k_a_), and dissociation constant (k_d_). We have immobilized WW1-WW2 FNBP4 on the CM5 sensor chip as a ligand, and different concentrations of FH1-FH2 FMN1 or FH2 FMN1 flowed over the immobilized surface as analytes in respective independent experiments. A significant increase in positive response was detected in the case of FH1-FH2 FMN1 as analyte (Fig. 2 B & C). FH1-FH2 FMN1 interacting with WW1-WW2 FNBP4 at high affinity, and K_D_ value was calculated as 1.84 nM, k_a_ .31*10^6^ M^-1^s^-1^, and k_d_ 5.705*10^-4^ s^-1^. When we used FH2 FMN1 as an analyte it did not interact with WW1-WW2 FNBP4 (Fig. 2 D). Finally, we examined WW1-WW2 FNBP4 binding activity with FH1-FMN1. From our data FH1-FMN1 construct expressed as 25 kDa protein despite FH1-FMN1 expected molecular weight is 14.4 kDa (Fig. 3 A & B). This FH1-FMN1 aberrantly migrate due to presence of poly-proline-rich sequences. Our SPR data confirmed that WW1-WW2 FNBP4 interacts with FH1 domain of FMN1 (Fig. 3 C). So, our results suggested that poly-proline-rich FH1 domain of FMN1 is responsible for binding with the WW1-WW2 FNBP4.

**Fig. 3.**
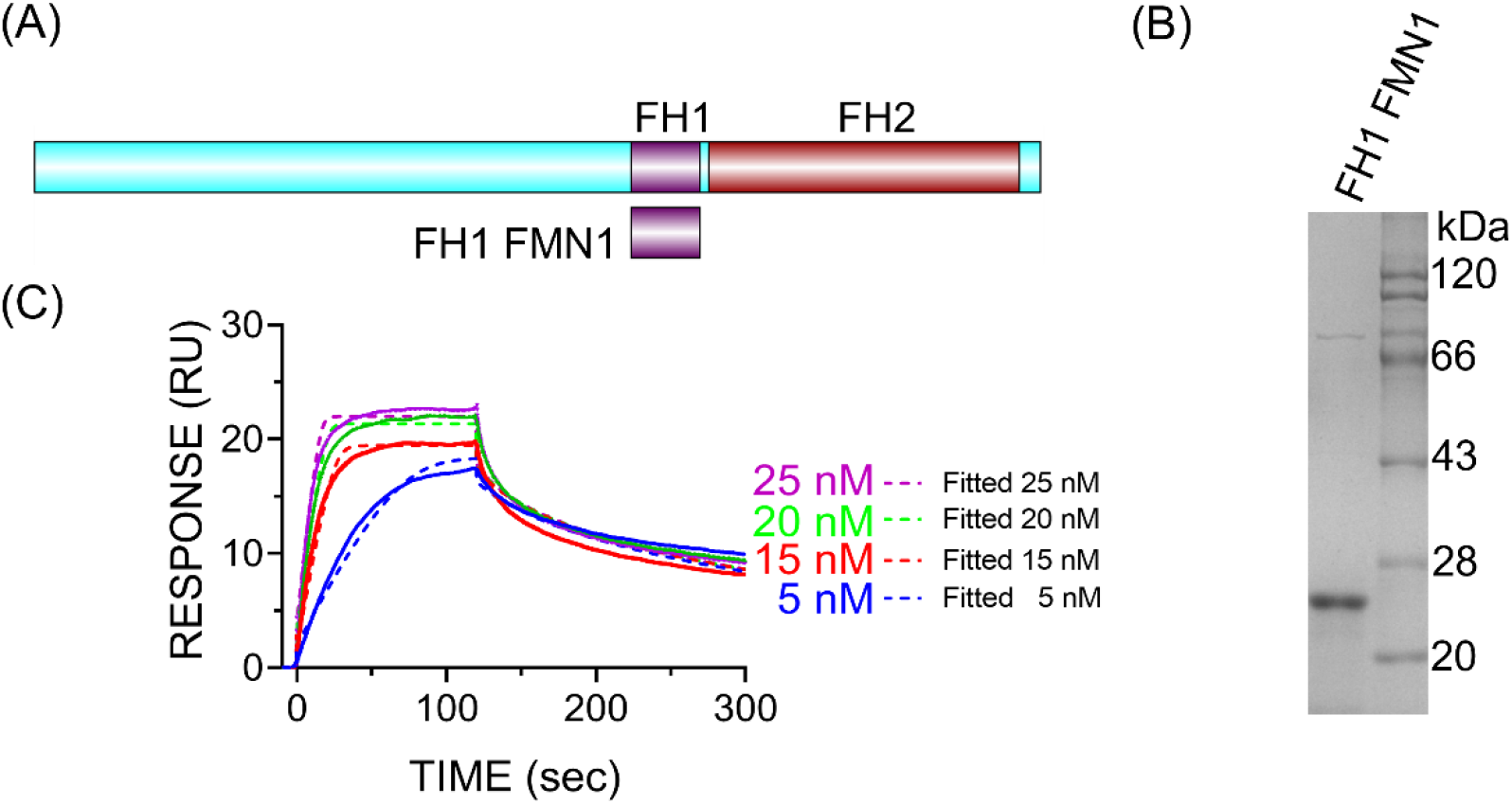
Binding between WW1-WW2 domain of FNBP4 and poly-proline rich FH1 domain of FMN1. (A) Schematic representation of FH1 FMN1 (amino acids 870-970) construct. (B) Coomassie stained 10% SDS-PAGE of purified 6x-His tagged FH1 FMN1 (amino acids 870-970). (C) SPR sensorgrams of the binding kinetics for WW1-WW2 FNBP4 with FH1 FMN1 at 25°C. WW1-WW2 FNBP4 was covalently immobilized on a CM5 (Series S) biosensor surface at pH 4.5 Increasing concentration of FH1-FMN1 (5 nM, 15 nM, 20 nM, and 25 nM) are shown in colored sensorgrams. Association phase was 120 sec and dissociation phase was 180 sec. Sensorgrams were fitted to Langumir binding rate equation and indicated with respective colored dash-line. Sensorgrams were blank-corrected for all cycles.

### 3.2. WW1 and WW2 domain resemblance in FNBP4

Human and mouse FNBP4 sequences were retrieved from GenBank (accession no. Q8N3X1 and Q6ZQ03 respectively). Multiple sequence alignment of WW1 (amino acids 214-248) and WW2 (amino acids 595-629) domain from FNBP4 had been checked using ClustalW. The sequence alignment showed WW1 domain is 31.43% identical to the WW2 domain (Fig. 4 A). It is noteworthy that the WW1 and WW2 domain of humans are fully identical to the WW1 and WW2 domain of mouse FNBP4 respectively (Fig. S. 3 A & B).

**Fig. 4.**
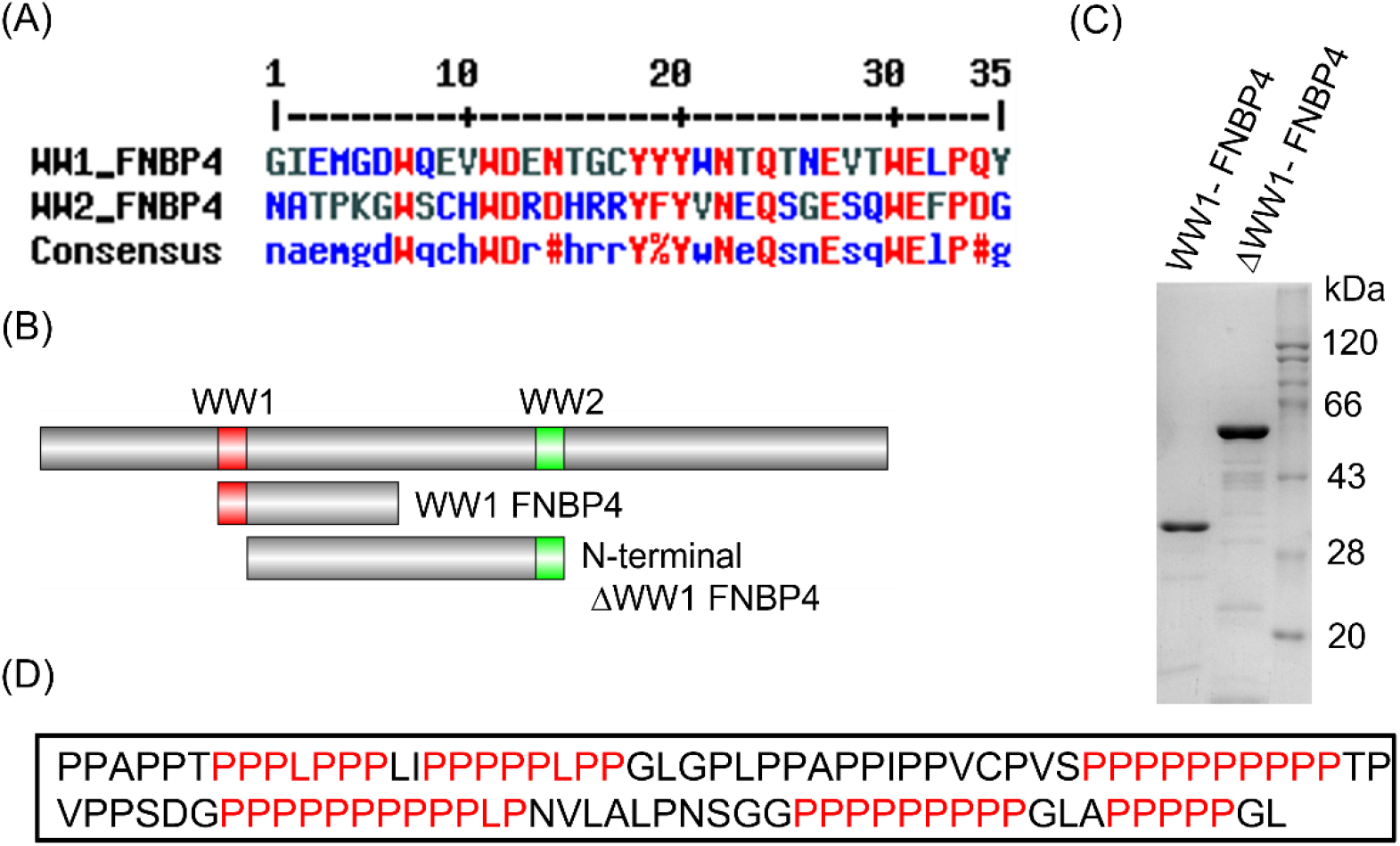
Sequence alignment, characteristic and purification of WW1 and WW2 domain of FNBP4. (A) Multiple sequence alignment of WW1 and WW2 domain of human FNBP4. Conserved amino acid residues were highlighted in red color. (B) Schematic representation of WW1 FNBP4 (amino acids 214-430) and N-terminal ΔWW1 FNBP4 (amino acids 249-629) constructs. (C) Coomassie stained 10% SDS-PAGE of purified 6x-His tagged WW1 FNBP4 (amino acids 214-430) and N-terminal ΔWW1 FNBP4 (amino acids 249-629) (26.7 kDa and 44.8 kDa from left to right). (D) Sequence of FH1 domain of FMN1 showing poly-proline consensus sequences, are highlighted in red.

### 3.3. WW1 and WW2 domain of FNBP4 behave differently

To distinguish the role between WW1 and WW2 domains of FNBP4, we compared the binding activity of N-terminal ΔWW1 FNBP4 and WW1 FNBP4 with FH1-FH2 FMN1. In ELISA, WW1 FNBP4 was coated in the well and incubated with different concentrations of FH1-FH2 FMN1 or FH2 FMN1 as analytes. We find FH1-FH2 FMN1 significantly binds to WW1 FNBP4 whereas FH2 FMN1 has no binding (Fig. 5 A). SPR data also corroborates the above result that WW1-FNBP4 binds with FH1-FH2 FMN1 (Fig. 5 B & C). Binding sensorgram revealed that WW1-FNBP4 fragment binds with FH1-FH2 FMN1 fragment with 2 nM K_D_ and in addition, their k_a_ and k_d_ value was 0.6372*10^6^ M^-1^s^-1^ and 1.273*10^-3^ s^-1^ respectively.

**Fig. 5.**
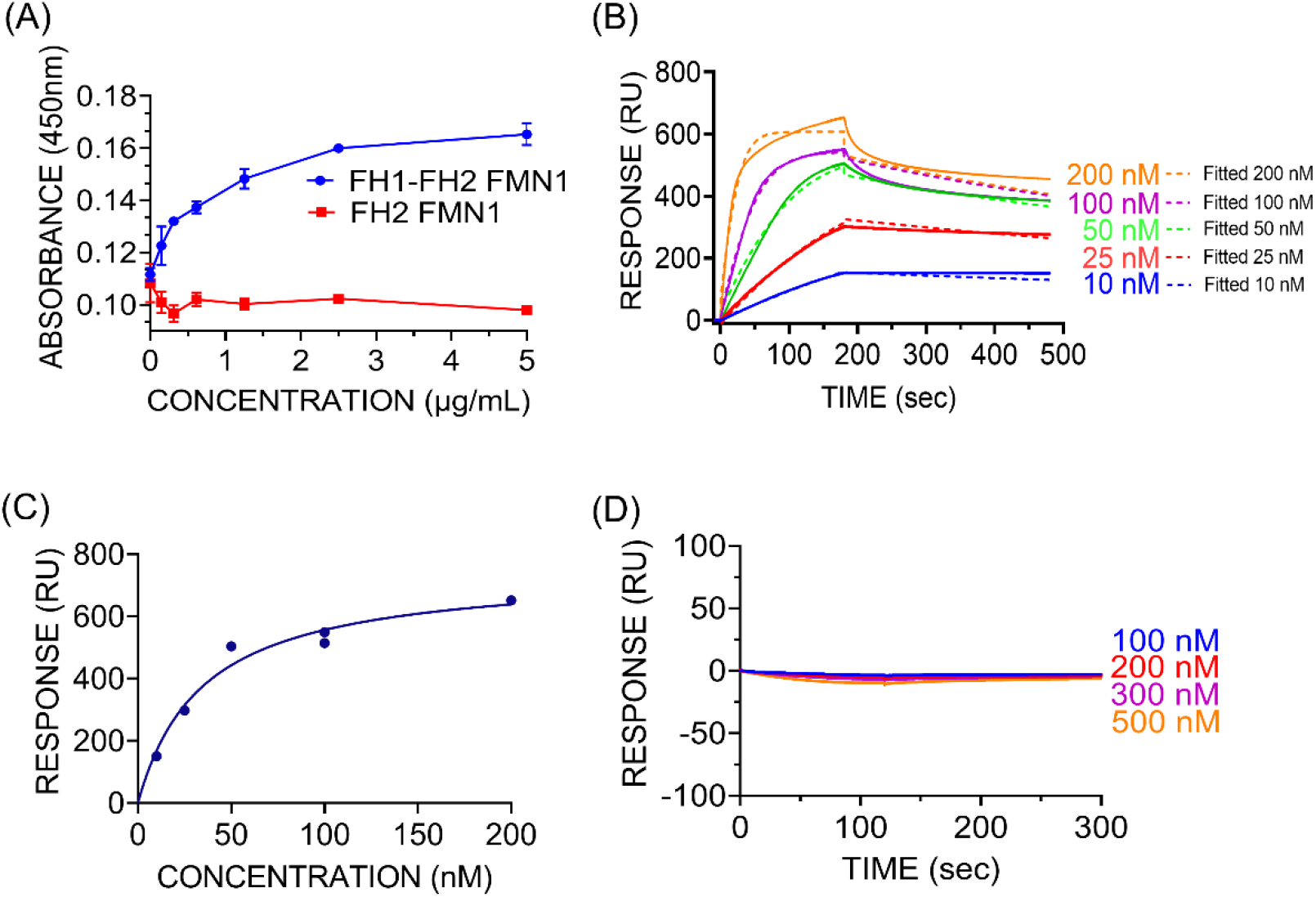
Binding between WW1 FNBP4 and poly-proline rich FH1 domain of FMN1. (A) Results of WW1 FNBP4 interaction with FH1-FH2 FMN1 and FH2 FMN1 obtained by ELISA. WW1 FNBP4 coated in well. FH1-FH2 FMN1 and FH2 FMN1 were used as analytes. (B) SPR sensorgrams of the binding kinetics for WW1 FNBP4 with FH1-FH2 FMN1 at 25°C. WW1 FNBP4 was covalently immobilized on a CM5 (Series S) biosensor surface at pH 4. Increasing concentration of FH1-FH2 FMN1 (10 nM, 25 nM, 50 nM, 100 nM & 200 nM) are shown in colored sensorgrams. Association phase was 180 sec and dissociation phase was 300 sec. The Langumir binding rate equation was used to fit the sensorograms, which are shown with the corresponding-colored dashed lines. (C) Binding affinity curve for FH1-FH2 FMN1 with WW1 FNBP4 was fitted with a non-linear regression equation (one-site specific binding model). Duplicate concentration of 100 nM was used for positive control (D) SPR sensorgrams of the binding kinetics for N-terminal ΔWW1 FNBP4 with FH1 FMN1 at 25°C. N-terminal ΔWW1 FNBP4 was covalently immobilized on a CM5 (Series S) biosensor surface at pH 4.5. Increasing concentration of FH1 FMN1 (100 nM, 200 nM, 300 nM, and 500 nM) are shown in colored sensorgrams.

In SPR analysis, we had immobilized ΔWW1 FNBP4 on the CM5 sensor chip and FH1-FMN1 flowed over the ligand immobilized sensor surface. The binding sensorgrams revealed that the ΔWW1 FNBP4 fragment did not bind to FH1-FMN1 (Fig. 5 D). We also examined N-terminal ΔWW1 FNBP4 binding activity with FH1-FH2 FMN1. From our SPR analysis, we had shown that N-terminal ΔWW1 FNBP4 did not interact with FH1-FMN1 (Fig. S. 4). So, our results had suggested that only the WW1 domain of FNBP4 responsible for binding with the poly-proline-rich FH1 domain of FMN1 where WW2 domain of FNBP4 did not involve in the interaction. Furthermore, we had also tried to dock FH1 domain of FMN1 and WW1 domain of FNBP4 using ClusPro2.0 (Fig. S. 5).

### 3.4. WW1 and WW2 domains of FNBP4 are independent

We checked the interaction between the WW1 and WW2 domains of FNBP4. WW2 FNBP4 construct was purified as 6x-His tagged with molecular weights of 29.9 kDa (Fig. S. 6 A & B). Then SPR analysis had been performed; where WW1-WW2 FNBP4 immobilized on CM5 chip, and WW1 FNBP4 and WW2 FNBP4 had flowed over the immobilized surface. It was observed in both cases that the sensorgrams generated negligible negative responses close to base line (Fig. 6 A &B). Above result indicates that WW1 does not interact with the WW2 domain of FNBP4.

**Fig. 6.**
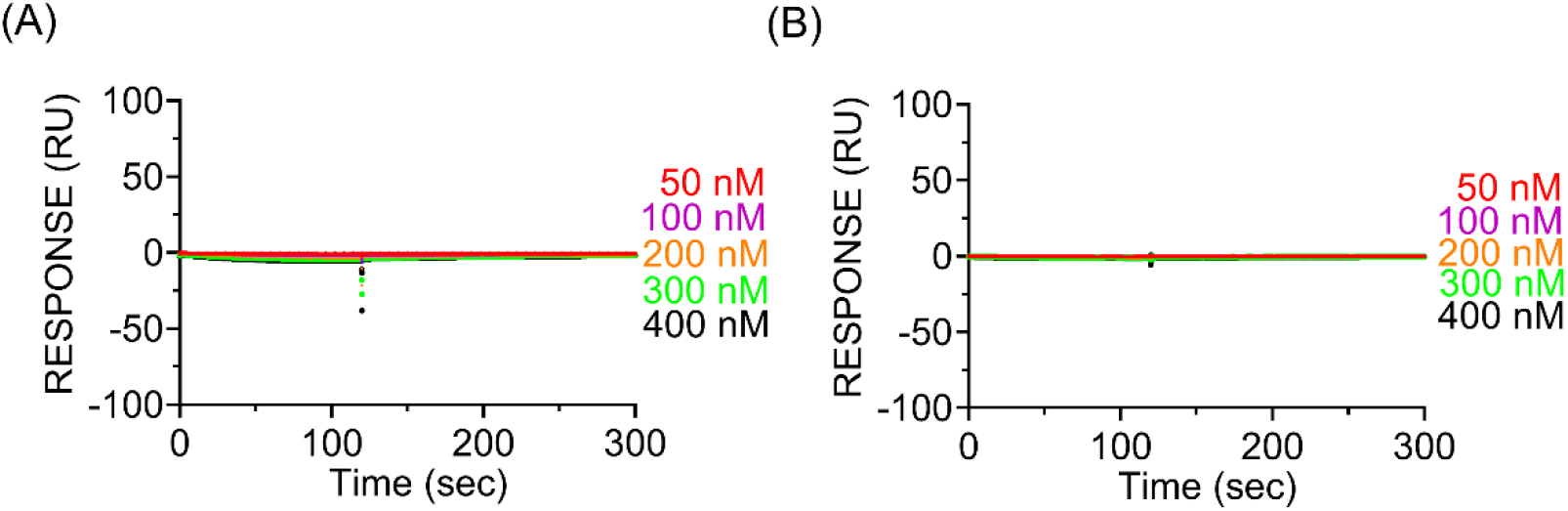
WW1 and WW2 does not exhibit interdomain interaction. SPR sensorgrams of the binding kinetics for WW1-WW2 FNBP4 to WW1 FNBP4 (A) and WW1-WW2 FNBP4 to WW2 FNBP4 (B) at 25°C. Various concentrations of WW1 FNBP4 or WW2 FNBP4 (50 – 400 nM) passed over WW1-WW2 FNBP4 immobilized surface.

Therefore, there was no inter-domain interaction between WW1 and WW2. In addition, we concluded that no intra-domain interactions exist since the WW1 FNBP4 and WW2 FNBP4 constructs are available to bind freely with WW1 and WW2 domains of the WW1-WW2 FNBP4 immobilized ligand, respectively.

## 4. DISCUSSION

In this study, we had deciphered the interaction between FNBP4 and FMN1 with high affinity. Presumably, the high affinity of FNBP4 for FMN1 might play a significant role in their functional regulations. So far FMN1 has not yet been characterized in terms of its regulation. FH1 domain of FMN1 contains several proline-rich consensus sequences (Fig. 4 D). Proteins like Fyn, Src, and FBPs protein afford to bind with poly-proline-rich stretches through their EVH1, SH3, and WW domains respectively.^25^ FH1 domain of Formins can interact with the SH3 domain-containing proteins. Remarkably the binding motif of the SH3 domain is PxxP, which is also similar to the PPxPP binding motif of WW domains.^15^ Additionally, SH3 and WW domains have similar XP2 binding grooves.^26^ Maria J Macias *et al*. showed bioinformatically SH3 domain of Sem5 protein and WW domain of Npw38 protein has a similar conserved binding site or binding pocket for ligand.^15^ As a consequence, it is possible to ideate that FBPs influence the binding of SH3 domain-containing proteins to interact with the FH1 domain.

In previous reports, proline-rich short synthetic peptides were used as ligands to examine their interaction with WW domains.^14,27^ Due to abundant poly-proline-rich sequence FH1 domain is an unstructured region. We had also tried to dock FMN1 and FNBP4 interaction, we feel that FH1 domain is unstructured when present in an unbound condition, but it might undergo structure formation upon binding with FNBP4 (Fig. S. 5).

FNBP4 contains two WW domains, where WW1 and WW2 are spaced by long stretches of amino acid. To address the different binding specificity of the WW1 and WW2 domains, we prepared WW1 and WW2 domain-containing constructs and checked their binding potentialities. From our results, it’s clear that the FNBP4 WW1 domain was only involved in interaction with the poly-proline-rich FH1 domain of FMN1. In contrast, WW2 domain did not bind to the FMN1 FH1 domain. Therefore, there was no binding, so ligand accessibility might not likely be the reason. WW2 domain of FNBP4 fine-tuned for different motifs that are not present in FMN1 FH1 domain. Different binding specificity of WW1 and WW2 might allow different formins to interact and play important role in cytoskeleton regulation.

In the endocytic pathway, the suppressor of Deltex [Su(dx)] interacts with the PY motif of Notch.^28^ Su(dx) has four WW domains (WW1, WW2, WW3, and WW4); individually or in pairs WW1-WW2 and WW3-WW4 can interact with Notch.^28^ Interestingly WW3 associates with the WW4 domain and obstruct WW4 to reach proper folding structure for Su(dx).^28^ FBP21 interacts with pre-mRNA splicing factor SIPP1 with the help of two tandem WW domains.^29^ FBP21 binding to SIPP1 was diminished when either one of the WW domains was mutated.^29^ This is clear evidence of the cooperation between the two WW domains in FBP21. In contrast, the tandem domains of FNBP4 might independently fold and behave differently for ligand binding (Fig. S. 7). Moreover, N-terminal WW1-WW2 FNBP4 together and WW1 FNBP4 solo confer a similar binding affinity for the FH1 domain of FMN1. From our results, it was clear that WW1 and WW2 did not interact with each other. Consequently, there was no co-operativity between WW1 and WW2 of FNBP4 for this FMN1 FH1 domain binding context or with other interacting partners also. Increasing corroboration assist the notion that in some tandem WW domain there is co-operatively due to the pliable linker region of the interdomain. FBP28, FBP11, and Su(dx) are all these protein’s tandem domains linked with a short flexible linker region.^30^ Whereas FNBP4 WW1 and WW2 domain were spaced by a long stretch around 347 amino acid residues might be the probable reason for solo acting rather than synergistically. However, the solo and tandem domains’ regulation and binding kinetics are very poorly understood. Extensive structural studies of the tandem FNBP4 WW domain will allow us to decipher the molecular picture of the regulation and co-operativity.

In previous reports, WW domains are classified as Gr-I, II, III, and IV based on binding with proline-rich consensus sequence. Group-I WW domains bind with PPxY motif, where x can be any acid residues including proline; Group-II recognizes PPLP consensus motif; Group-III prefers to bind with poly-proline stretch flanked by an arginine amino acid at C-terminus, and Group-IV showed an affinity for phospho-Serine/Threonine proceeded by proline amino acid residues.^27^ Later due to ambiguity in the grouping of Gr-II and III as both Gr-II and Gr-III domains can bind to PPLP and PPR motifs along with simple poly-proline stretch, Gr-II/III is considered a single group.^26^ FNBP4 WW domains were categorized in this large group-III based on prior studies utilizing short proline peptides containing PPR consensus sequences.^27^ Here, we observed that the FNBP4 WW-1 domain fulfills these Gr-II / III criteria for the motif binding preference. Nevertheless, the FNBP4 WW-2 domain was classified into Gr-II / III, but further study using long ligands is needed to correctly classify this domain.

Finally, FNBP4 interacting partners of FMN1 have been identified, since FNBP4 and FMN1 both have been reported as a regulator of the BMP signaling pathway for limb development. The expression pattern of FNBP4 and FMN1 in different developmental phases in the mice model should shed additional light on this. FNBP4 is a multi-domain protein that might form multiprotein complex with FMN1 formin and other protein, where N terminal of FNBP4 interact with FMN1 and C-terminal domain of FNBP4 might interact with other protein. Future research is required to determine if the C-terminal domain of FNBP4 interacts with additional proteins.

## Declarations

### Conflicts of interest/Competing interests

The authors declare no competing interests.

### Author contributions

SM; project conceptualization, designing, funding acquisition, manuscript writing-review and editing.SD; designing, conducting biochemical experiments, analysis, and manuscript writing.

## Acknowledgements

SM thanks DST-FIST for funding the Central Analytical Instrumentation Facility and IISER-KOLKATA for extending institutional funding.SD acknowledges the University Grants Commission for fellowship. We thank Mr. Somnath Halder at Central Analytical Instrumentation Facility at IISER-Kolkata for his assistance during SPR experiments. A special thanks to Mr. Jajati Keshari Ray for his assistance with the animal house facilities. We thank Prof. S Gourinath and Dr. Pragyan Parimita Rath for helping in the molecular docking studies. We are grateful to Prof. Rupak Datta and Dr. Arnab Gupta for their valuable comments in the preparation of the manuscript.

## Supplementary figure, and legends

**Fig. S 1.**
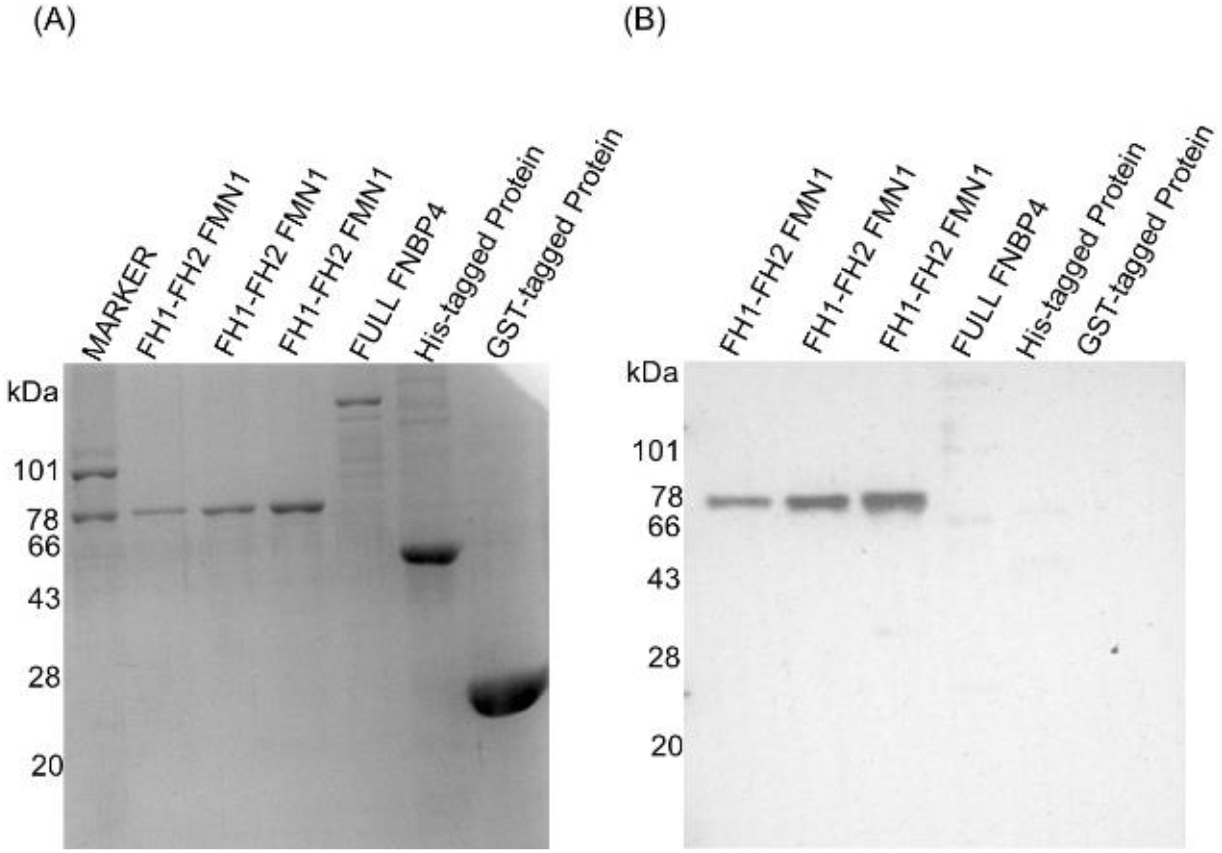
Raised FH1-FH2 FMN1 Antibody is specific for FMN1 recombinant proteins. (A) Coomassie-stained 10% SDS-PAGE replica gel showing different recombinant protein samples. (B) Reactivity of the FH1-FH2 FMN1 constructs with our raised anti-FMN1 sera by Western blot analysis.

**Fig. S 2.**
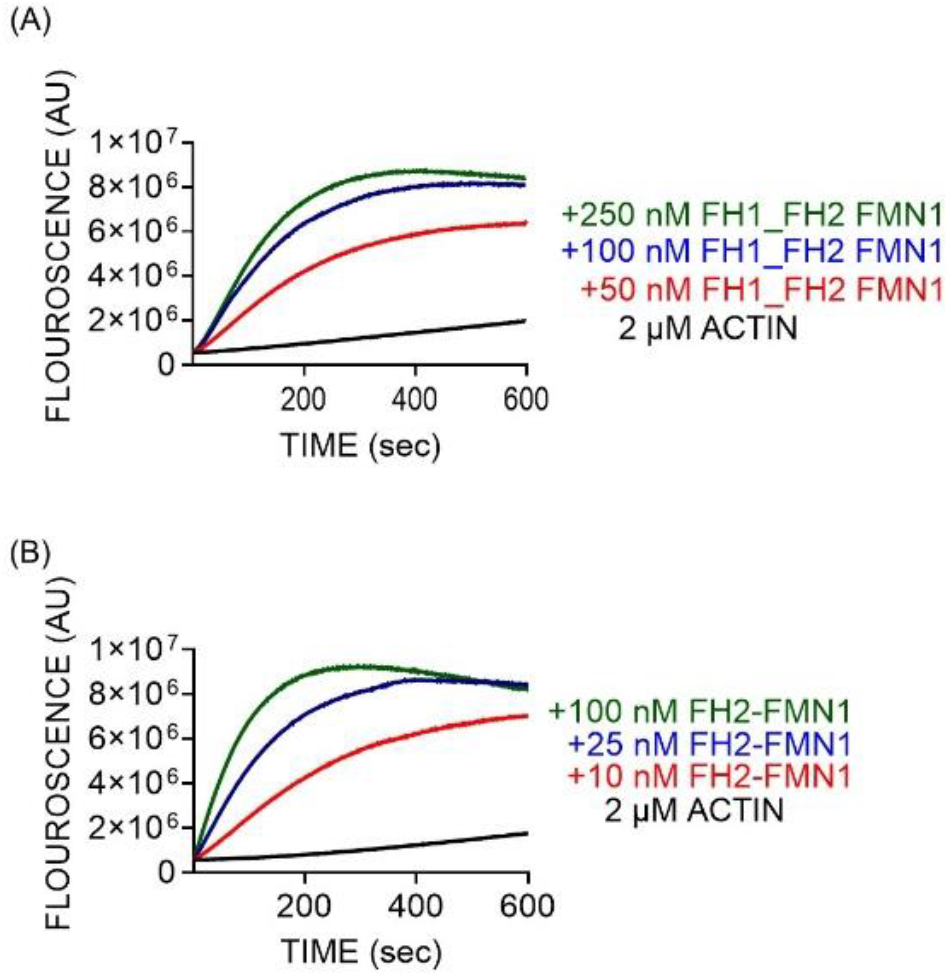
Effect of FH1-FH2 FMN1 or FH2 FMN1 on spontaneous actin polymerization. 2 μM 10% pyrene-labeled actin polymerization assay in presence of (A) FH1-FH2 FMN1, and (B) FH2 FMN. Different Color-coded lines indicate increasing concentrations of analytes. Pyrene labeled actin concentration was kept at 2 μM. Pyrene fluorescence was monitored at 365 nm excitation and 407 nm emission spectra.

**Fig. S 3.**
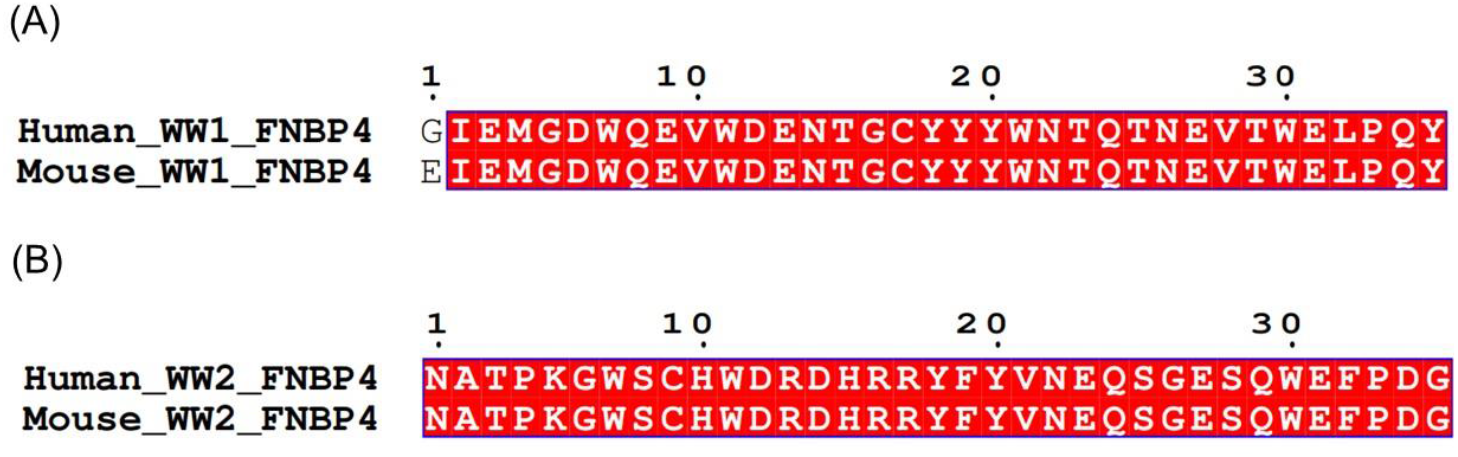
Multiple sequence alignment of WW1 and WW2 domain of FNBP4 from human and mouse. (A) Human WW1 domain and mouse WW1 domain (B) Human WW2 domain and mouse WW2 domain. Conserved amino acid residues were highlighted in red color.

**Fig. S 4.**
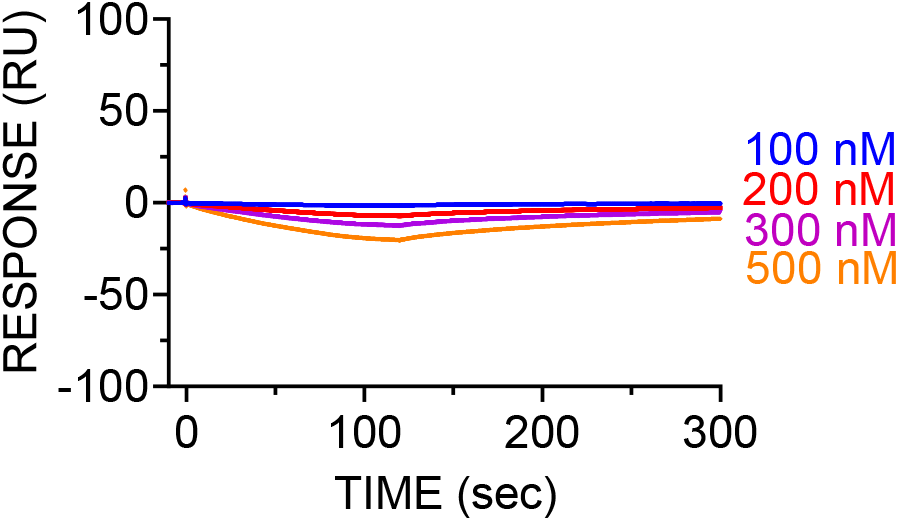
ΔWW1 FNBP4 does not interact with poly-proline rich FH1-FH2 domain of FMN1. SPR sensorgrams of the binding kinetics for N-terminal ΔWW1 FNBP4 with FH1-FH2 FMN1 at 25°C. N-terminal ΔWW1 FNBP4 was covalently immobilized on a CM5 (Series S) biosensor surface at pH 4.5. Increasing concentration of FH1-FH2 FMN1 (100 nM, 200 nM, 300 nM, and 500 nM) are shown in colored sensorgrams.

**Fig. S 5.**
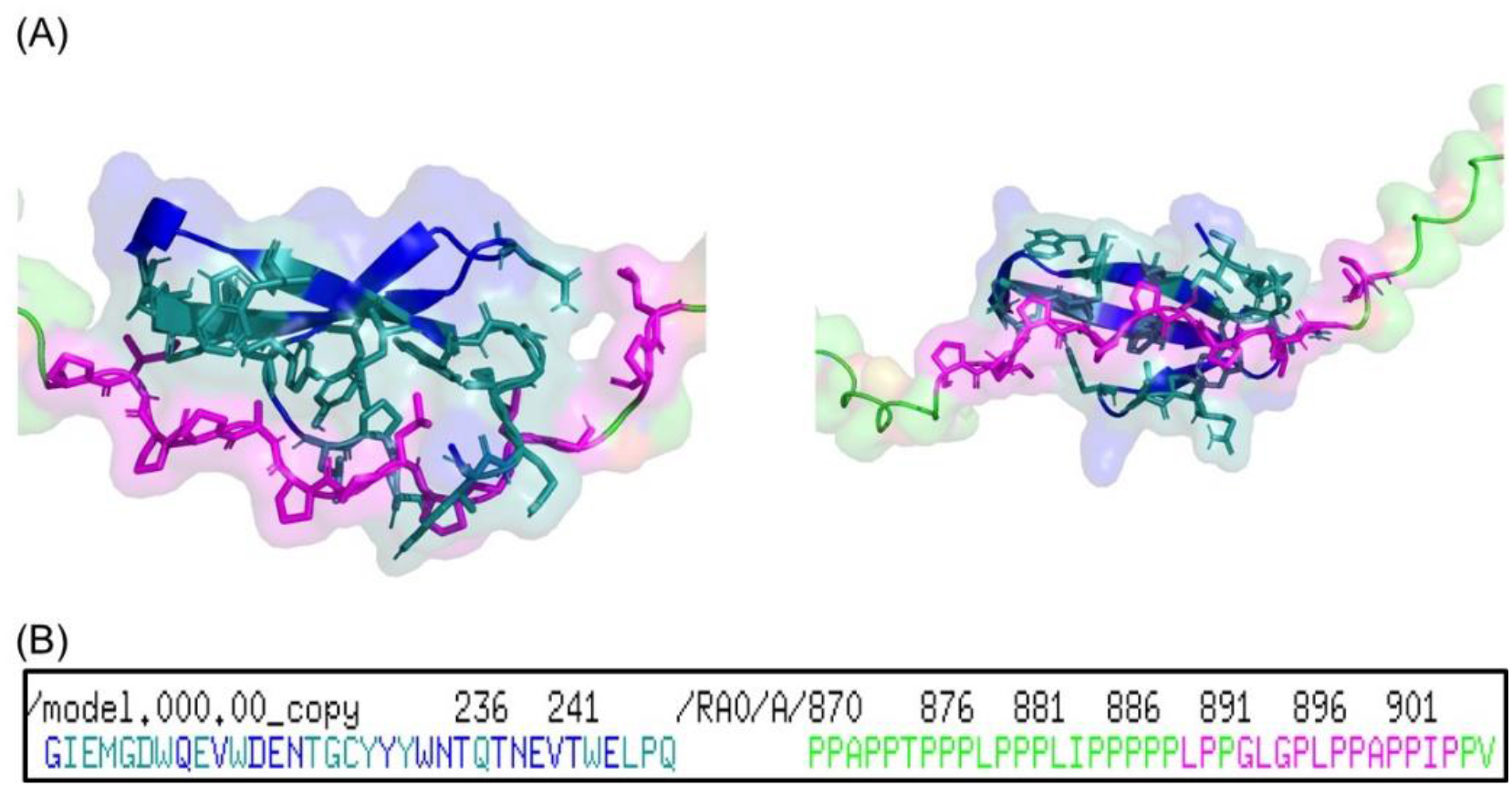
Results of molecular docking between FH1 domain of FMN1 and WW1 domain of FNBP4. (A) Blue coloured structure is WW1 region of FNBP4 protein, while Green coloured structure is FH1 region of the formin protein. The Teal coloured residues from WW1 protein and Magenta residues from FH1 region represent the interacting residues in the docked structure. (B) Predicted interactions were displayed in the figure, two protein regions were colored separately for clearer visualization.

**Fig. S 6.**
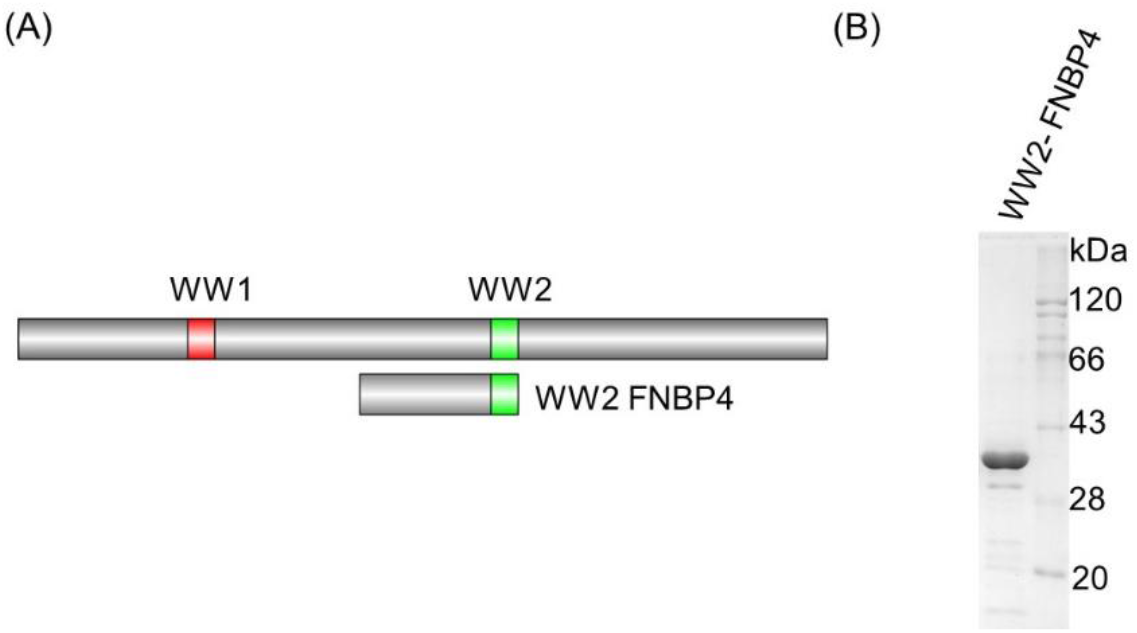
Schematics of construct and purified fragment of FNBP4. (A) Schematic representation of WW2 FNBP4 (amino acids 384-629) construct. (B) Purified WW2 FNBP4 (amino acids 384-629) in coomassie stained 10% SDS-PAGE (30 kDa).

**Fig. S 7.**
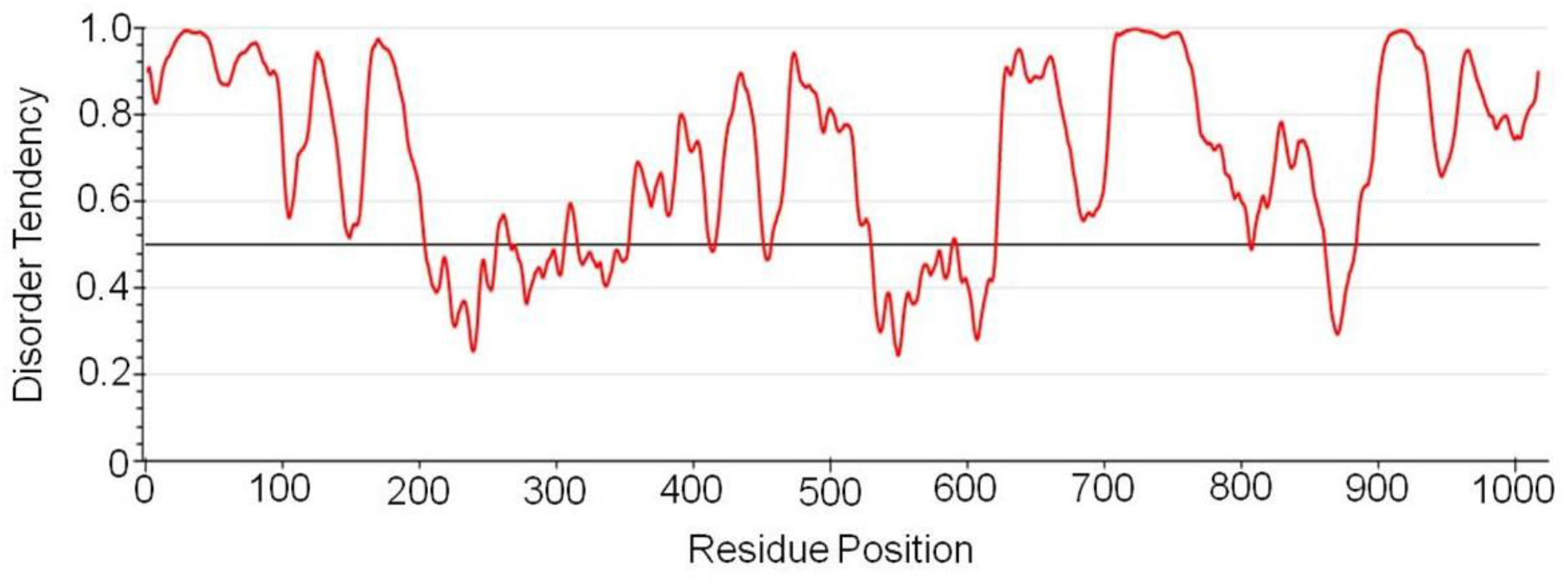
IUPred3 Disorder prediction graph for FNBP4, where values greater 0.5 indicates a tendency to be disordered region.

## Supplementary material, and methods

**Table 1:**
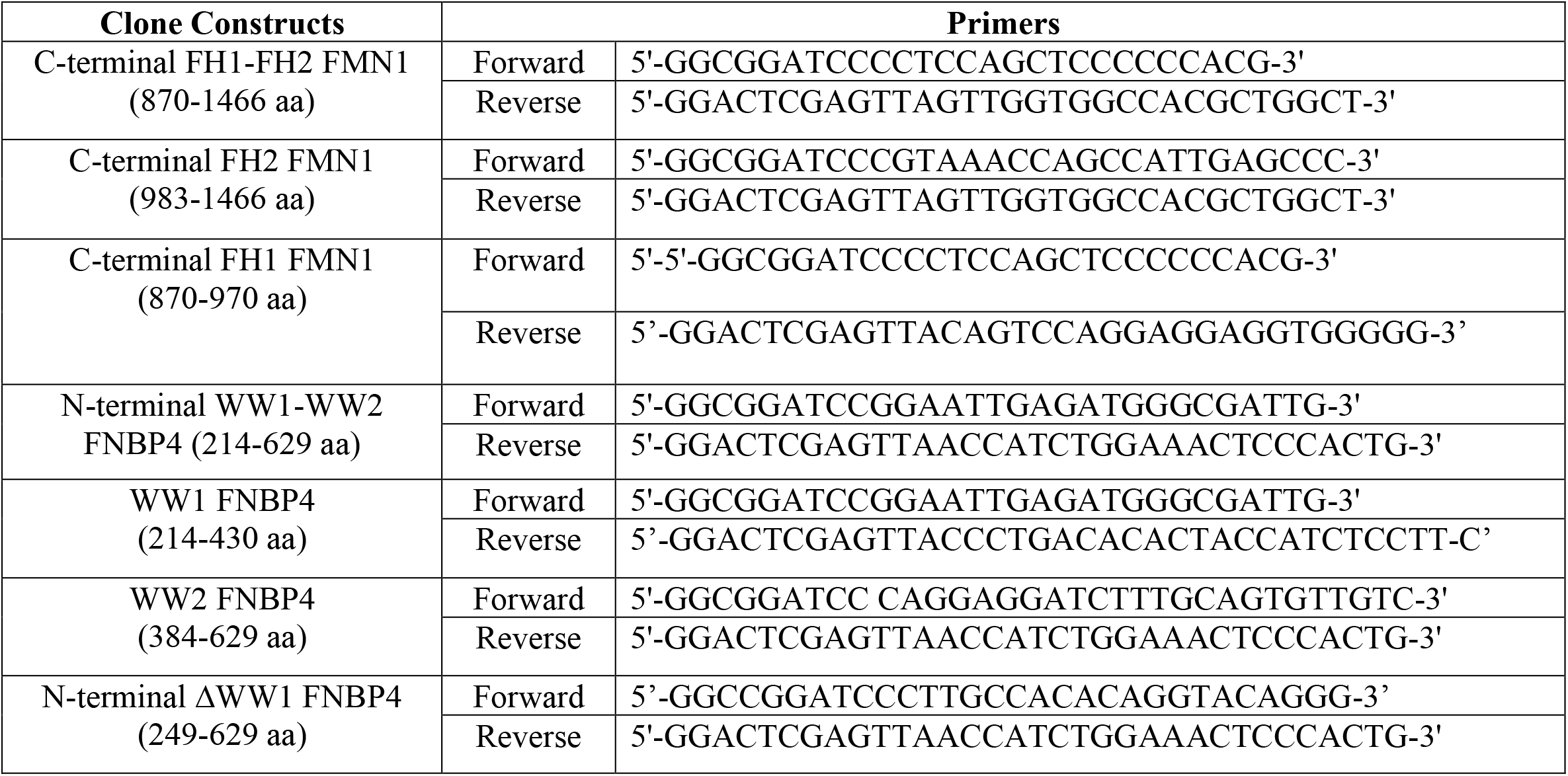
List of primers used for cloning of FMN1 and FNBP4 constructs.

### Pyrene–actin polymerization assays

The purification of G-actin was carried out using rabbit muscle acetone powder^1^. N-(1-pyrene) iodoacetamide was used to label rabbit muscle actin.^2^ At first, I had added 27.8μL of G-buffer (10 mM Tris at pH 7.5, 0.2 mM ATP, 0.2 mM DTT, 0.2 mM CaCl_2_). Then I had added 2μM (10 μL) of G-actin from the 12 μM (10% labeled G-actin and 90% unlabeled G-actin) stock solution. Then 4.2μL of exchange buffer (10 mM EGTA, 1 mM MgCl_2_) was added to the solution and incubated for 2 minutes at room temperature. Next, the candidate protein (FH1-FH2 FMN1 or FH2 FMN1) was added in different concentrations and the final volume was adjusted with HEK buffer (20 mM HEPES at pH 7.5, 1 mM EGTA, 50 mM KCl, 5% glycerol). Then actin polymerization was initiated with the addition of 3 μL 20X Initiation Mixture (1 M KCl, 40 mM MgCl_2_, and 10 mM ATP). The final reaction volume was 60μL. Reaction was monitored at 365 nm (excitation spectra) and 407nm (emission spectra) in a fluorescence spectrophotometer (Photon Technology International).^3^

### Immunoblotting

Protein samples were run in 10% SDS-PAGE and blotted onto the PVDF membrane (#IPVH00010, Merck). A 5% skimmed milk solution was used to block the PVDF membrane for two hours at room temperature. Primary antibody incubation is performed overnight at 4°C, followed by HRP-tagged secondary antibody incubation at room temperature for 2 hours. To obtain the chemiluminescence signal, the manufacturer’s protocol was followed using SuperSignal West Pico Chemiluminescent Substrate (#34080, Thermo Scientific).

### Protein-protein docking

We downloaded the Alphafold2 models for both FMN1 and FNBP4 proteins. Next, we used the WW1 region from FNBP4 and FH1 region from Formin1 for docking studies using ClusPro2.0. The model with the highest Lowest Energy weighted score (−612.6) from the ClusPro2.0 docking was chosen for analysis. As displayed in the figure, the predicted interaction was found between the two protein regions (colored separately for clearer visualization).

### Protein disorder prediction

We have predicted the structural disordered region of FNBP4 using IUPred3 (https://iupred3.elte.hu/) bioinformatics server. The IUPred3 graph for FNBP4, where greater values equate to a higher probability of disorder (unstructured), was used to determine the disorder propensity of each residue in the FNBP4.

## Notes

### Competing Interest Statement

The authors have declared no competing interest.

